# DIA-PASEF Proteomic Profiling Reveals MpkA-Dependent Iron Stress Responses and Siderophore Biosynthesis in *Aspergillus nidulans*

**DOI:** 10.1101/2025.07.03.662993

**Authors:** JungHun Lee, Olivia K. West, Walker D. Huso, Alexander G. Doan, Kelsey J. Grey, Harley Edwards, Jasmine T. Tran, Dylan R. Carman, Michael J. Betenbaugh, Ranjan Srivastava, Steven D. Harris, Mark R. Marten

**Author notes:** Harley Edwards is currently at 1601 Research Boulevard Rockville, MD 20850, United States. **Corresponding Author** Corresponding author at: Engineering, Room 314, 1000 Hilltop Circle, Baltimore, MD 21250, United States. E-mail address (M.R. Marten).

## Abstract

Filamentous faungi play essential roles in biotechnology as producers of valuable bioproducts and, conversely, as opportunistic pathogens. *Aspergillus nidulans* is a widely used model organism for fungal genetics and cell biology; however, comprehensive proteomic references for this species remain limited. In this study, we applied a data-independent acquisition–parallel accumulation serial fragmentation (DIA-PASEF) approach to enable efficient and in-depth proteome profiling of *A. nidulans*. Leveraging ion mobility-based ion cloud information from DDA-PASEF experiments, we developed DIA-PASEF methods that identified 3,904 proteins across biological triplicates grown in rich medium. Compared to prior studies, this represents an increase in protein identifications of more than 140% and was achieved with more than five-fold reduction in analysis time. We employed this newly developed DIA-PASEF methodology to conduct both proteomic and phosphoproteomic analyses under iron-depleted conditions in an MpkA protein-kinase deficient mutant (Δ*mpkA*). The Δ*mpkA* strain exhibited expression of approximately 500 additional proteins and occupancy of over 1,800 additional phosphosites relative to a control. Differentially expressed and phosphorylated proteins increased by more than an order of magnitude in the Δ*mpkA* mutant across both iron-replete and iron-deplete conditions. Gene Ontology (GO) enrichment analysis revealed broader and distinct biological processes under iron-depleted conditions, highlighting adaptive responses specific to iron limitation and MAPK pathway disruption. This work establishes a high-coverage proteomic resource for *A. nidulans* and provides novel insights into fungal stress responses and signaling network perturbation. Importantly, high-throughput proteomic profiling reveals that limited iron availability and MAPK pathway disruption increases siderophore biosynthesis.

## Introduction

Filamentous fungi are important in the fields of both biotechnology and medicine, serving as prolific producers of bioactive compounds and, conversely, as opportunistic pathogens ^1, 2^. To optimize beneficial applications and improve treatment strategies, a comprehensive understanding of fungal systems biology is essential. The species *Aspergillus nidulans* is widely used as model system to study fungal cell biology. Proteomic approaches, in particular, enable high-throughput insights into fungal protein expression and regulation ^3–5^. High-resolution mass spectrometry approaches, such as quadrupole time-of-flight (QTOF) and orbitrap, serve as a powerful platforms for shotgun proteomics ^6–8^. Yet, prior analyses have often yielded limited proteome coverage, requiring extensive gradients or multiple sample fractions ^5, 9–11^. Trapped ion mobility time of flight spectrometry (timsTOF) improves ion separation and scan efficiency, and when coupled with PASEF (Parallel Accumulation–Serial Fragmentation), enables rapid and sensitive peptide analysis ^12, 13^. Thus, the timsTOF system offers the potential of both deeper and faster proteomic profiling.

Numerous investigations into the fungal proteome have predominantly relied on data-dependent acquisition (DDA) methodologies ^5, 9, 11, 14–16^. DDA offers the advantage of efficiently profiling large peptide populations in a relatively short analysis time ^17^. Nonetheless, its inherent bias toward selecting high-abundance precursor ions limits its capacity to detect low-abundance proteins. In contrast, data-independent acquisition (DIA) addresses this limitation by systematically fragmenting all detectable precursor ions at the MS1 level, and capturing them at MS2, thereby enhancing proteome coverage and enabling the identification of a broader proteome range ^18^. However, the increased data complexity of DIA can lead to prolonged computational times during peptide identification. DIA-PASEF, an integrated technology combining DIA and PASEF, leverages ion mobility separation to reduce scan times ^13^. By targeting uniquely charged precursor ion clouds based on ion mobility and m/z, DIA-PASEF enables faster and more comprehensive proteomic analyses ^18–20^.

Iron serves as a critical cofactor in numerous biological processes due to its ability to alternate between ferrous and ferric states ^21^. During microbial infection, host cells often utilize iron restriction as a defense mechanism in an attempt to limit pathogen proliferation ^21, 22^. Thus, to establish infection, pathogens must overcome this limitation by competing with the host for access to essential iron resources. Microbial pathogens have been reported to utilize siderophore-mediated systems for efficient iron acquisition, a mechanism recognized as an important virulence factor. ^23^.

In a previous study ^4^, we found iron homeostasis in the model fungus *Aspergillus nidulans* was impacted by deletion of the protein kinase MpkA. And others have shown similar results in the pathogen *Aspergillus fumigatus* ^4, 24^. While MpkA is a crucial component of the cell wall integrity (CWI) signaling pathway, which regulates cell wall remodeling and repair in response to environmental stress ^4, 5, 25^, it is not completely clear how this is related to iron-related metabolism. Thus, investigating proteome alterations in the model organism *Aspergillus nidulans*, under iron- and MpkA-deficient conditions can provide insights regarding the molecular mechanisms underlying fungal response to iron deficiency and resultant pathogenicity.

With these ideas in mind, we used a proteomic analysis approach to assess alterations in protein expression and phosphorylation in response to iron-deficient conditions in an *mpkA* deletion mutant. We employed a DIA-PASEF approach to achieve deeper proteome coverage (compared to DDA) of *Aspergillus nidulans* while maintaining short acquisition times. To enhance peptide identification efficiency and accuracy, we first generated a spectral library using high-pH fractionation. Compared to previous studies, our workflow yielded significantly more protein identifications, in a shorter timeframe, from a small number of samples.

We used our developed DIA-PASEF workflow to investigate the impact of MpkA protein kinase on the *A. nidulans* whole- and phosphoproteome under both iron-sufficient and iron-depleted conditions, revealing their combined influence on siderophore synthesis

## Methods

### Strains and Media

*Aspergillus nidulans* strains A1405 (control) and A1404 (Δ*mpkA* derivative of A1405) were obtained from the Fungal Genetics Stock Center (Manhattan, KS, USA) ^4, 5^. We have reported genotypic details previously ^25^. Strains were pre-cultured on MAGV-S agar (20 g/L malt extract, 20 g/L agar, 20 g/L glucose, 2 g/L peptone, 325.2 g/L sucrose, 0.1% (v/v) Hunter’s trace elements, and 0.1% (v/v) vitamin solution) at 28 °C for three days. For DDA-PASEF and DIA-PASEF protein identification and library generation, 10⁷ spores of A1405 were inoculated into triplicate low-pH YGV medium (pH 3.3) in 250ml flasks and incubated for 12 hr, followed by transfer to triplicate 1.2 L YGV medium (pH 6.0) in 2.8L Fernbach flaks and cultured for 20 hr at 28 °C, 250 rpm, for library expansion 20 ng micafungin/ml was added at 20hr and collected after 2 hr, as previously described by Chelius et al. (2019) and Chelius et al. (2020) ^4, 5^. For iron and MpkA deficiency studies, triplicate seed cultures were grown in modified minimal medium (pH 3.3) as described by Chelius et al. (2019) for 16 hr and transferred to iron-replete or iron-depleted medium (with/without FeSO₄) for an additional 20 hr before harvesting. Iron-deficient culture flasks were pre-treated with 2 M HCl for 24 hr to remove trace metals ^4^.

### Protein digestion

Mycelia were collected with Miracloth filtration, frozen in liquid nitrogen, and ground to powder using mortar and pestle under frozen conditions. Proteins were extracted in TNE buffer (20 mM Tris-HCl, 150 mM NaCl, 2 mM EDTA) and quantified by BCA assay (Pierce, Rockford, IL, USA). Protein, 500 μg, was precipitated using cold 20% TCA, washed with cold acetone, and digested using the Filter-Aided Sample Preparation (FASP) method as described by Chelius et al. (2019) ^4,26^. Samples were reduced with TCEP, alkylated with iodoacetamide, washed with NH₄HCO₃, in 3kDa ultrafiltration unit (MilliporeSigma, Burlington, MA, USA) and digested with trypsin (1:50 enzyme:sample) overnight at 37 °C. Digestion was quenched with trifluoroacetic acid (TFA) to final concentration of 0.1%.

### Desalting and phospho-enrichment

Solid-phase extraction was used for peptide desalting with a 1cc Oasis HLB cartridge (Waters, Milford, USA) ^27^. The cartridge was conditioned by acetonitrile containing 0.1% formic acid, followed by equilibration with 0.1% formic acid. Peptide samples were loaded onto the cartridge and washed three times with the equilibration buffer. Elution was performed using a stepwise gradient of acetonitrile (40%, 60%, 80%, and 100%) containing 0.1% formic acid, then dried in SpeedVac. Phosphopeptides were enriched using the High-Select™ TiO₂ Phosphopeptide Enrichment Kit (Thermo Scientific), following the manufacturer’s instructions and the method described by Chelius et al. (2019) ^4, 28^.

### Fractionation

High-pH reverse-phase fractionation was performed using the A1405 YGV peptide sample ^29^. Oasis HLB cartridge was conditioned and equilibrated with acetonitrile followed and 15 mM ammonium formate buffer (pH 10.0) ^27, 30^. The desalted peptide sample was loaded onto the cartridge, then washed with equilibration buffer. Peptides were eluted using a stepwise gradient of acetonitrile containing 15 mM ammonium formate (pH 10.0) at 20 different concentrations (2, 5, 7.5, 10, 12.5, 15, 17.5, 20, 22.5, 25, 27.5, 30, 32.5, 35, 40, 50, 60, 70, 80, and 100%). The elutes were dried immediately then, subsequently pooled into eight fractions.

### LC/MS analysis

Peptides were resuspended in 2% acetonitrile with 0.1% formic acid, then analyzed on a nanoElute LC system (Bruker, Billerica, MA, USA) coupled to timsTOF Pro2 mass spectrometer (Bruker, Billerica, MA, USA) at the Molecular Characterization and Analysis Complex (MCAC), University of Maryland, Baltimore County (UMBC). Samples, 1 μg, were separated on a PepSep C18 column (Bruker, Billerica, MA, USA) (15 cm × 75 μm, 1.9 μm particle size) at 50 °C using 18- and 150-minute gradients from 5% to 30% acetonitrile in 0.1% formic acid, followed by a rapid increase to 95%. MS scan was performed in positive ion mode using both DDA-PASEF and DIA-PASEF acquisition. TIMS ramp times were set to 100 ms. For DDA-PASEF, precursors were selected with an intensity threshold of 5,000, and 4 or 10 MS/MS scans were acquired per cycle (18- or 150-min gradients). Fragmentation was performed with collision energies ramping from 20.00-65.00 eV in the range of 0.60-1.60 Vs/cm² (18 min) or 17.00-75.00 eV for 0.50-1.78 Vs/cm² (150 min). For DIA-PASEF 18-min gradient, 8 MS/MS ramps and 14-27 MS/MS windows were used with −0.95 s cycle time. Ion mobility and charge state information from 2+ DDA-PASEF spectra were used to guide DIA window design.

### Peptide search

DDA-PASEF data were searched using MSFragger (version 4.1) against the *A. nidulans* UniProt database (UP000000560) ^31^. Variable modifications included oxidation (M) and N-terminal acetylation, and phosphorylation (S/T/Y only for phospho-proteome), with carbamidomethylation (C) as fixed. Mass tolerance was set to 20 ppm; up to three variable modifications and two missed cleavages were allowed. FDR was controlled at 1%. DIA data were analyzed using DIA-NN (version 1.9.1) with the same search parameters, applied to the DDA-PASEF data ^32^. The spectral library generated from DDA-PASEF analysis was used as input for DIA-PASEF processing. Statistical Analysis and Gene Ontology Annotation Protein group data from DIA-NN were log₂-transformed (LFQ intensities) and analyzed in Perseus (version 2.1.3.0) ^33^. Two-sample t-tests were performed (p < 0.05). Proteins with fold-change >2 or <0.5 and p < 0.05 were defined as significantly differentially expressed or phosphorylated. String-db (https://string-db.org/) was used for Gene Ontology (GO) enrichment ^34^.

### Siderophore Assay

Seed cultures of *A. nidulans* strains A1405 and A1404 were prepared by inoculating 10⁷ spores into 50 mL of modified minimal medium (pH 3.3) and incubating for 16 hours. Subsequently, 2.1 ml of seed culture was transferred to 50 ml of minimal medium (pH 6.5) and cultured for an additional 20 hours. For iron-depleted conditions, FeSO₄ was omitted from the minimal medium, and culture flasks were pre-treated with 2 M HCl for 24 hours to eliminate residual trace metals. Dry cell weight (DCW) was measured, and 2 mL of culture supernatant was filtered through a 0.22 μm membrane for siderophore quantification. Siderophore levels were determined using the Chrome Azurol S (CAS) assay as described by Chelius et al. (2019). Briefly, 100 μl of filtered supernatant was mixed with 100 μl of CAS reagent, and absorbance at 630 nm was measured after 10 minutes. Siderophore production (%) was calculated using the formula:

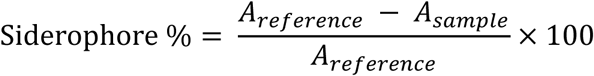

where *A_reference_* and *A_sample_* represent the absorbance of the control and sample, respectively. Siderophore production was normalized to biomass by dividing the siderophore percentage value by the corresponding dry cell weight. Statistical significance was assessed using a t-test, and all analyses were performed using IBM SPSS Statistics (version 30.0.0.0) software.

## Results and Discussion

### Expansion of *A. nidulans* proteome identification

To generate a comprehensive spectral library, we grew the *A. nidulans* control strain (A1405) in YGV medium for 20 h, followed by protein extraction and peptide preparation using high-pH reverse-phase fractionation ^4, 26, 29^. Peptides were collected in 20 fractions and pooled into eight to minimize overlap (**Figure 1A and Supplementary Table S1**). A spectral library was generated using DDA-PASEF data obtained in this study, including high-pH fractionated peptide proteome data. The DIA-PASEF method was then developed using ion mobility data from 2+ precursors, segmented into 14 acquisition windows (59.0 Da width) (**Figure 1A**). Proteomic profiling of the control strain (A1405) cultured in YGV with an 18-minute gradient yielded an average of 3,207 proteins by DDA-PASEF and 3,826 by DIA-PASEF, reflecting a 19.3% increase (**Figure 1B and Supplementary Table S2**). In total, 3,904 proteins were identified across triplicates by DIA-PASEF. Compared to prior studies, this reflects a >140% increase in protein identifications with over five-fold reduction in scan time ^5, 9–11^. Considering the 10,561 annotated proteins in *A. nidulans* (UniProt), this approach covered approximately 36% of the predicted proteome, demonstrating the utility of DIA-PASEF for high-throughput fungal proteomics.

**Figure 1.**
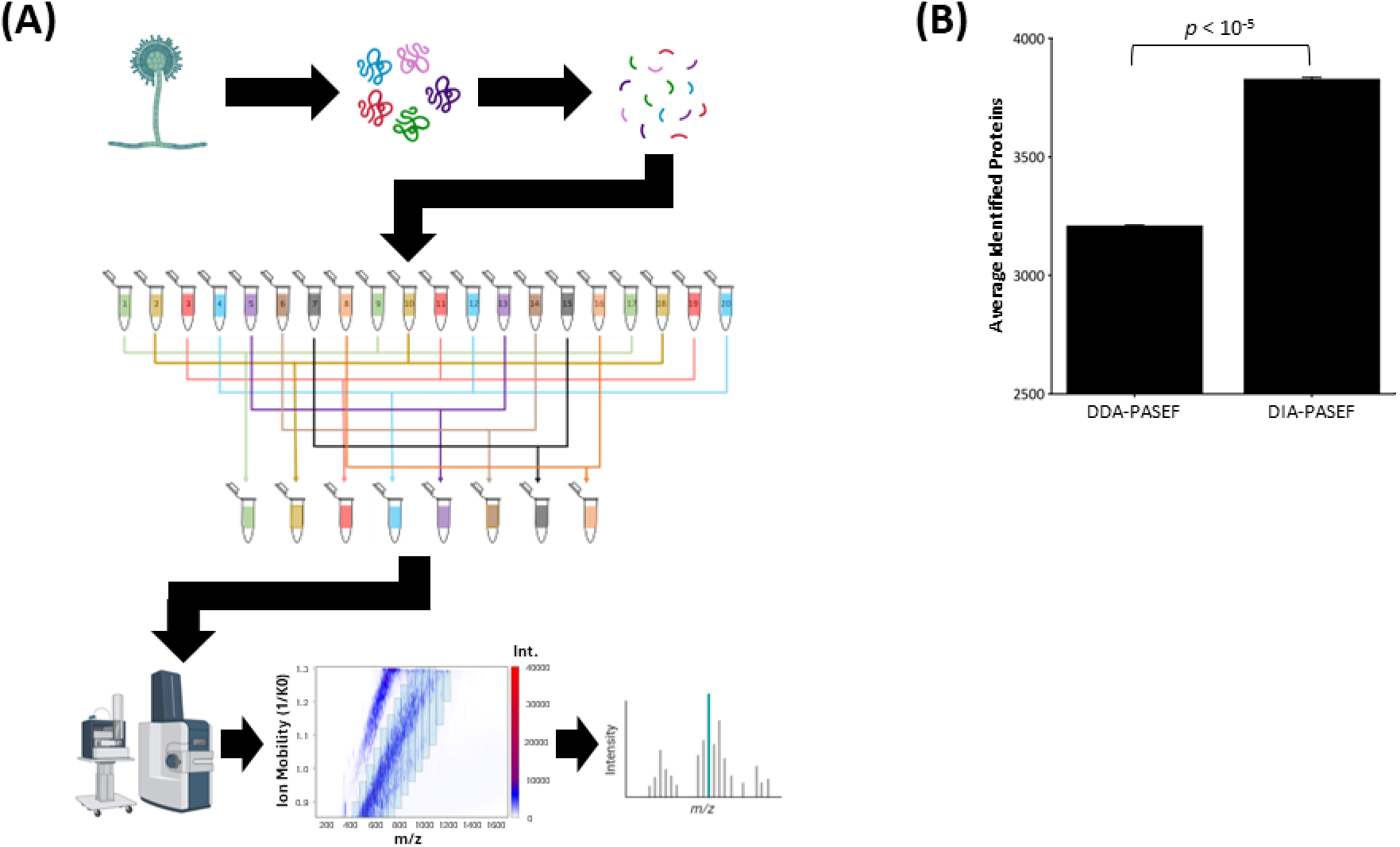
**(A).** Schematic overview of *A. nidulans* peptide fractionation for spectral library generation and DIA-PASEF analysis. **(B).** Average number of proteins identified using 18-minute gradient DDA-PASEF and DIA-PASEF methods.

### Impact of iron and MpkA deficiency in whole-proteome expression

In a previous investigation ^4^, we found that in the absence of cell-wall stress, deletion of the *A. nidulans* wall-repair gene, *mpkA*, leads to altered physiological characteristics including changes in iron homeostasis. To assess the combined effects of iron limitation and *mpkA* deletion, control (A1405) and Δ*mpkA* (A1404) strains were grown under iron-replete and iron-deficient conditions. DIA-PASEF whole-proteome profiling revealed an average of 3,446 and 3,422 proteins for control in iron-replete and iron-deficient media, respectively. In contrast, A1404 showed an average of 3,951 and 3,921 proteins under the same conditions (**Figure 2A and Supplementary Table S3**). Across all groups, 4,335 proteins were identified, with 3,428 proteins (∼80%) overlapping. Strain-specific comparisons revealed 117 and 555 unique proteins in A1405 and A1404, respectively, under iron-replete conditions; under iron deficiency, 191 and 571 unique proteins were observed (**Figure 2B**). Under iron-sufficient and iron-deficient conditions, A1405 exhibited 134 and 111 unique proteins, respectively, while A1404 showed 118 and 37 (**Figure 2B**). Condition-specific comparisons showed moderate shifts in protein identity, suggesting that *mpkA* deletion drives broader proteome remodeling independent of iron status. Statistical analysis identified differentially expressed proteins (DEPs) using cutoffs of *p* <0.05 and fold change >2 or <0.5. Volcano plots revealed 70 DEPs in A1405 across iron conditions, while only 16 were identified in A1404. Comparisons between strains under the same iron condition revealed more dramatic changes: 898 DEPs in iron-replete and 1,097 DEPs in iron-deficient conditions. These results again indicate that *mpkA* deletion has a more pronounced impact on global protein expression than iron availability alone (**Figure 2C**).

**Figure 2.**
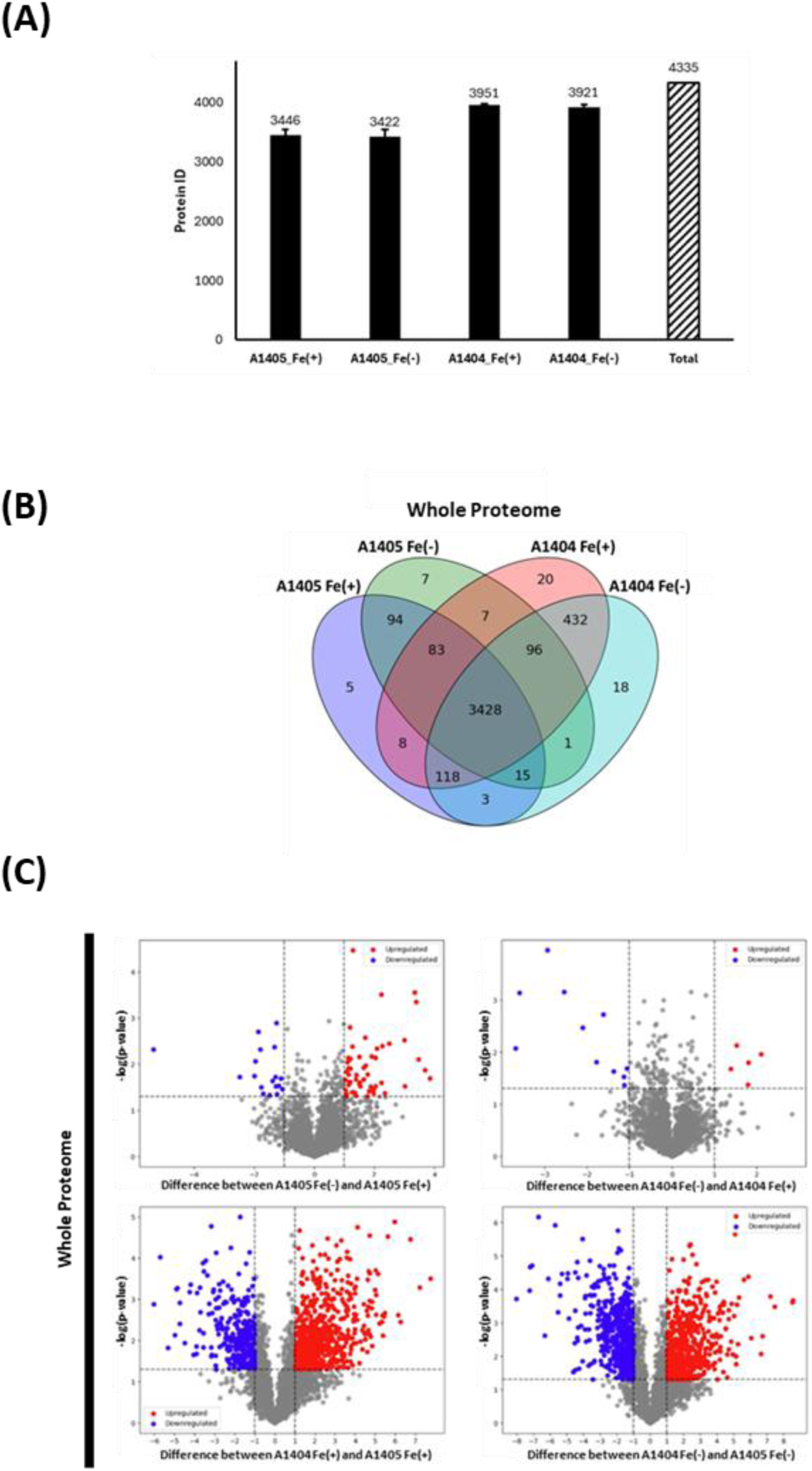
**(A).** Average number of proteins identified under iron and MpkA deficient conditions and total number of identified proteins. **(B).** Venn diagrams showing the overlap of identified proteins across cases. **(C).** Volcano plots illustrating differentially expressed proteins for the following comparisons (clockwise): A1405 Fe(-) vs Fe(+); A1404 Fe(-) vs Fe(+); A1404 Fe(+) vs A1405 Fe(+); and A1404 Fe(-) vs A1405 Fe(+).

### Impact of iron and MpkA deficiency in phosphoproteome expression

Given MpkA’s role in the cell wall integrity (CWI) signaling pathway, which operates through a phosphorylation mediated cascade ^4, 5, 25, 35–37^, we analyzed phosphorylation profiles as well. Using phospho-enriched peptides from the same samples, DIA-PASEF identified 4,421 phosphorylation sites across 1,700 proteins (**Figure 3A and Supplementary Table S3**). Despite similar total protein counts, A1404 exhibited an average of 3,023 and 3,096 phospho-sites in iron replete and deplete conditions, while A1405 showed an average of 1,102 and 1,295 phospho-sites, indicating a broader phosphorylation landscape in the mutant (**Figure 3A**). Strain-specific comparisons revealed 157 and 2,471 unique phosphosites in A1405 and A1404, respectively, under iron-replete conditions; under iron deficiency, 255 and 2,416 unique phosphosites were observed (**Figure 3B**). Under iron-sufficient and iron-deficient conditions, A1405 exhibited 170 and 352 unique proteins, respectively, while A1404 showed 332 and 361 (**Figure 3B**). Differentially phosphorylated proteins (DPPs) were defined using the same statistical cutoffs as DEPs. In A1405, 121 phospho-sites differed between iron conditions; A1404 showed 76. Strain comparisons revealed 1,858 DPPs under iron-replete and 2,103 under iron-deficient conditions (**Figure 3C**), emphasizing the role of MpkA in regulating phosphorylation, particularly under iron stress.

**Figure 3.**
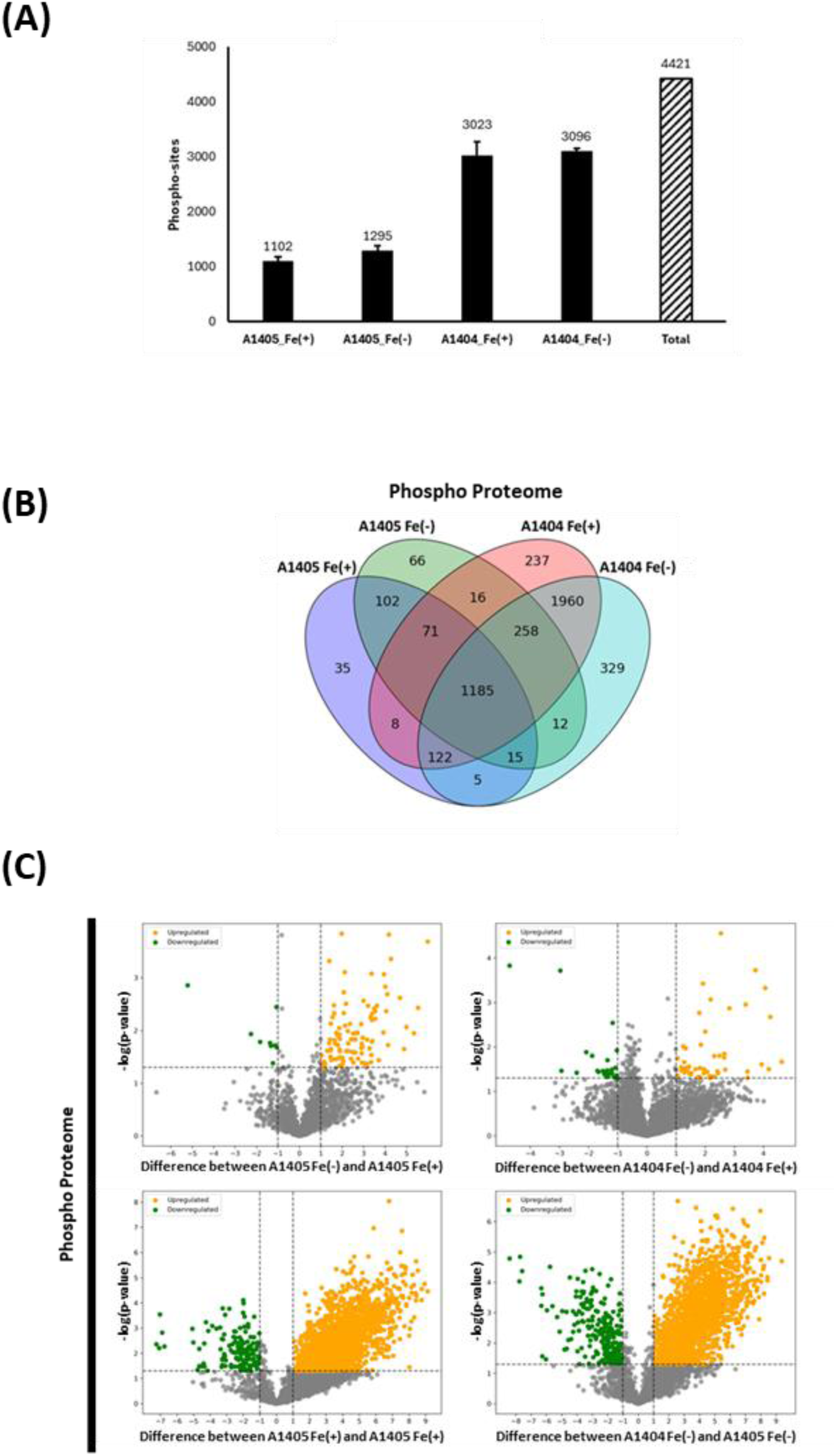
**(A).** Average number of phosphorylated proteins under iron and MpkA deficient conditions and total number of phosphorylated proteins **(B).** Venn diagrams showing the overlap of phosphorylated proteins across cases. **(C).** Volcano plots illustrating differentially phosphorylated proteins for the following comparisons (clockwise): A1405 Fe(-) vs Fe(+); A1404 Fe(-) vs Fe(+); A1404 Fe(+) vs A1405 Fe(+); and A1404 Fe(-) vs A1405 Fe(+).

### Gene Set Enrichment

GO enrichment analysis was performed to investigate the biological relevance of DEPs and DPPs (**Figure 4 and Supplementary Table S4**). Under iron-replete conditions, A1404 vs A1405 DEPs were enriched in stress response, amino acid metabolism, nucleotide metabolism, macromolecule metabolism, peptide metabolism, small molecule metabolism, and ribosome biogenesis. These categories expanded under iron deficiency, with additional unique enrichment in ATP/ADP metabolism and phosphate-containing compound metabolism (**Figure 4A and B**). For DPPs of A1404 vs A1405, enrichment under iron-replete conditions included cytoskeleton organization, macromolecule metabolism, phosphorylation, stress response, amino acid metabolism, and ribosome biogenesis. Under iron-deficient conditions, these categories were broadened with additional unique enrichment in transport and secretion, reproduction, and phosphorus metabolism (**Figure 4C and D**). To further investigate the impact of iron deficiency in the MpkA-deficient strain, we compared the protein expression and phosphorylation profiles of A1404 and A1405. These differences were uniquely observed and significantly enriched in the GO analysis under iron-deficient conditions (**Figure 5 and Supplementary Table S5**). All proteins associated with the ATP/ADP metabolic process were upregulated. Unique GO terms in A1404 under iron deprivation included “ATP/ADP metabolic process” and “Phosphate-containing compound metabolic process” (**Figure 5A**). Additionally, genes involved in the “Purine metabolism” and “Small molecule metabolic process” were significantly more represented in A1404 under iron-deficient conditions (**Figure 5A**). All the genes were upregulated with glycolytic enzymes such as *enoA* and *pgkA* ^38–40^. The overexpression of *adk2* suggests an active ATP/ADP exchange reaction in A1404 under deplete iron condition ^41^. These suggest increased nucleotide turnover. Also, MAPK pathway components, *sskB*, *hog1*, *ypdA*, *nimX*, AN3310.2, and AN4189.2, were overexpressed within the “Phosphate-containing compound metabolic process” category, highlighting potential cross-regulation between iron response and kinase signaling ^25, 42–44^. Additionally, the differential regulation of *AN3499.2*, *AN6037.2*, *AN5908.2*, *AN2334.2*, *AN2180.2*, *AN3169.2*, *AN1965.2*, *AN3223.2*, *enoA*, and *pgkA* implies that the pentose phosphate pathway (PPP) was upregulated, possibly to synthesize amino acids and nucleotides ^38–40, 45–51^. Positive regulation of the PPP, in response to iron limitation in A1404, was also indicated within the “Purine metabolism” category, presenting upregulation of *AN6708.2*, *ANIA_05655* (*AN5655*), *ANIA_06993* (*AN6993*), *ANIA_3466* (*AN3466*), and *AN1965.2*, while *AN6037.2*, *AN2334.2*, *AN5908.2*, *AN2180.2*, and *AN6815.2* downregulation ^43, 45, 47, 49, 50, 52–55^. Under the Gene Ontology term “Secondary molecule metabolic process,” several genes associated with the tricarboxylic acid (TCA) cycle were observed to be overexpressed, including *ANIA_03466* (*AN3466*), *ANIA_03639* (AN3639), *ANIA_05790* (AN5790), and *acuE*. The TCA cycle provides carbon skeletons for the biosynthesis of various amino acids, such as glutamate, arginine, and ornithine, key precursors for siderophore production in *Aspergillus nidulans* ^56–58^. Phosphorylation changes were uniquely observed in “Reproduction” GO category, including chitin synthases: ChsB (Q00757) and ChsC (P30583) were positively phosphorylated, while ChsD (P78611) was negatively phosphorylated in A1404 under iron deficiency (**Figure 5B**) ^59–61^. Meiosis-associated proteins, Q5AXD2, Q5B224, Q5B2W5, C8VG99, C8VQ48, and Q00083, showed enhanced phosphorylation, as did the asexual reproduction transcription factor P36011 ^62–68^. In the “Transport and Secretion” category, proteins such as C8VTT1, Q5AZS0, and Q5BFX0 were positively phosphorylated, while C8VTZ0 (Ras GTPase-related) and Q5AWT0 (Na⁺/H⁺ exchanger) were negatively modified ^69–72^. “Cytoskeleton” related proteins including C8V1M3, O74689, and Q5BBF2 exhibited increased phosphorylation in A1404 under iron deficiency, as did the actin cytoskeleton regulator Q5BEN1 ^73–77^. Under the “amino acid” category, two iron-binding related proteins, Q5B394 and Q5AVX6, were found to be differentially phosphorylated.^78, 79^. Overall, these results suggest that iron limitation in the MpkA deficient strain, A1404, leads to extensive remodeling of metabolic, reproductive, transport, and cytoskeletal pathways, with phosphorylation playing a significant regulatory role.

**Figure 4.**
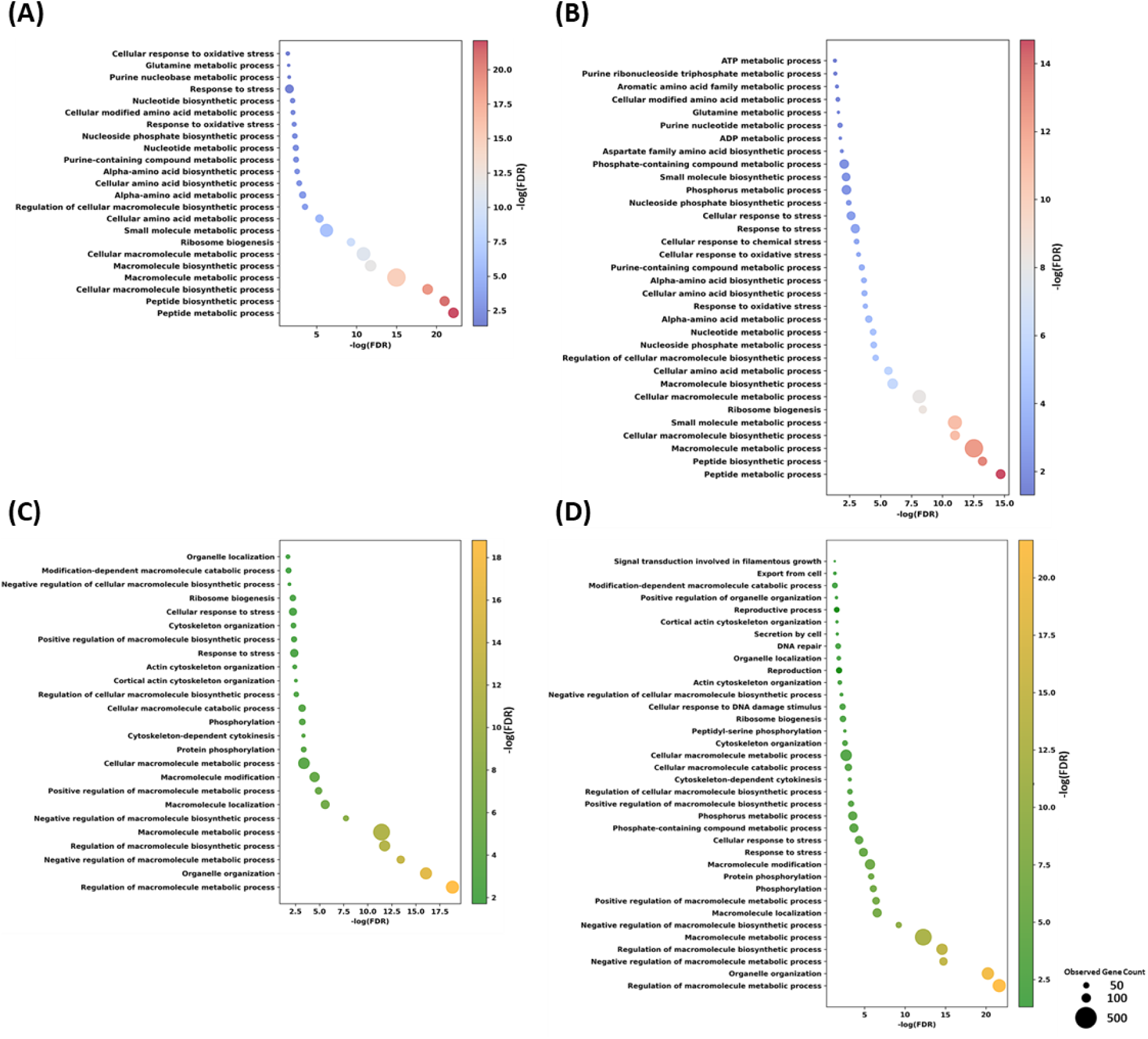
Dot plot of GO analysis of DEPs and DPPs. **(A)** DEPs in A1404 Fe(-) vs A1405 Fe(+). **(B)** DEPs in A1404 Fe(-) vs A1405 Fe(-). **(C)** DPPs in A1404 Fe(+) vs A1405 Fe(+). **(D)** DPPs in A1404 Fe(-) vs A1405 Fe(-).

**Figure 5.**
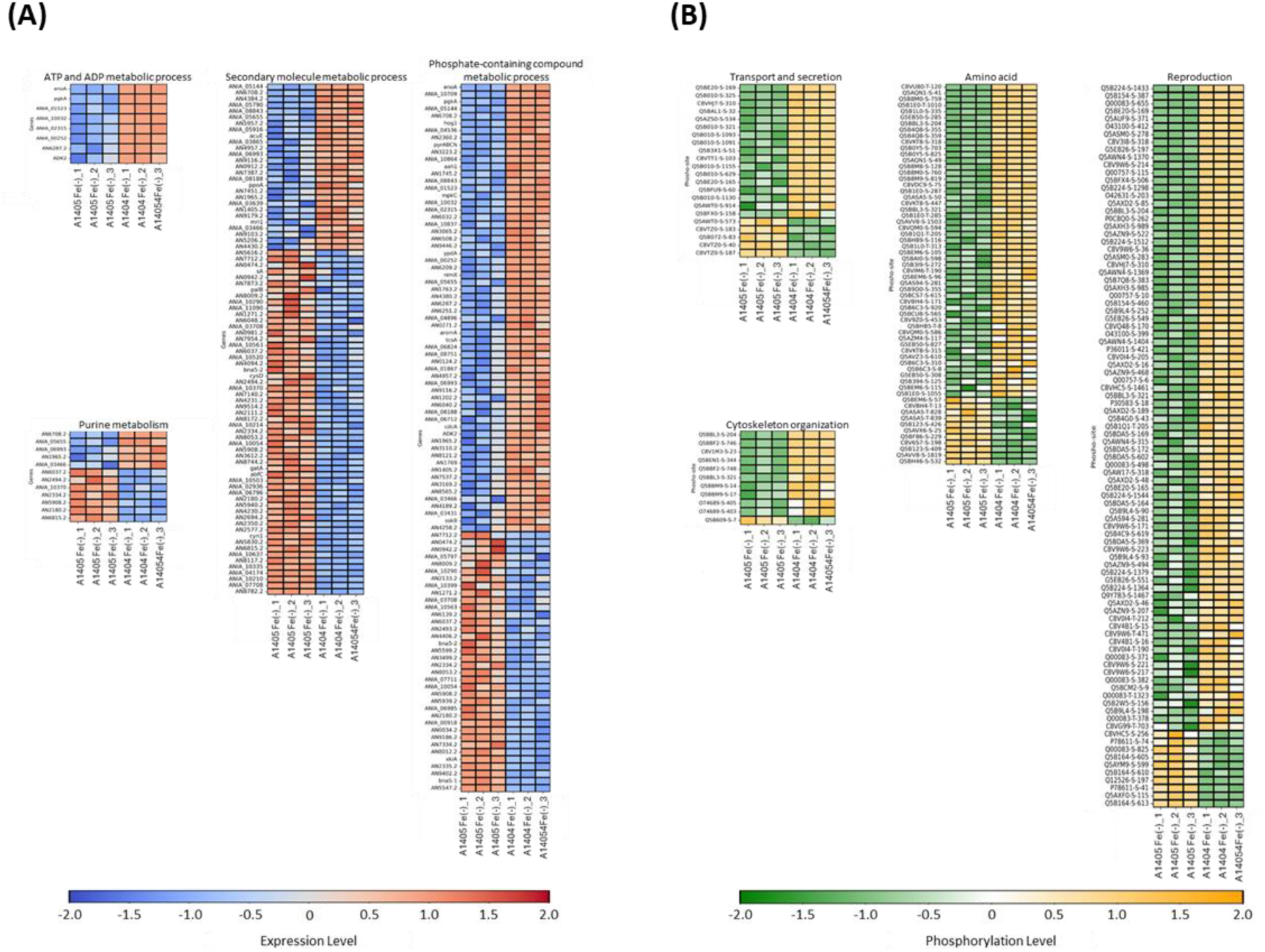
**(A)**. Heatmap of categorized genes exclusively expressed in A1404 Fe(-) vs A1405 Fe(-) comparison to A1404 Fe(+) vs A1405 Fe(+). **(B).** Heatmap of categorized genes exclusively phosphorylated in A1404 Fe(-) vs A1405 Fe(-) comparison to A1404 Fe(+) vs A1405 Fe(+).

### Siderophore production

Alterations in iron and MpkA conditions are known to influence siderophore biosynthesis ^4, 24^. Based on our GO enrichment analysis, we hypothesize that combined iron limitation and MpkA deficiency may lead to an increased ATP/ADP pool and enhanced carbon flux through the pentose phosphate pathway (PPP) and tricarboxylic acid (TCA) cycle. This metabolic reprogramming could contribute to elevated siderophore production. Specifically, upregulation of the TCA cycle may enhance the synthesis of key siderophore precursors, while an increased ATP/ADP pool may support the energy-intensive processes involved in siderophore biosynthesis ^41, 56–58^. To evaluate the impact of iron availability and MpkA on siderophore production in *Aspergillus nidulans*, we performed the Chrome Azurol S (CAS) assay. Compared to the control strain (A1405) under iron-sufficient conditions, the *ΔmpkA* (A1405) under iron-deficient conditions produced 2.89-fold more siderophore per unit biomass (**Figure 6**). Furthermore, strain A1404 under iron-deficient conditions exhibited a 32% increase in siderophore production per unit biomass relative to A1405 under the same iron-deficient condition (**Figure 6**). CAS assay performed on the A1404 Fe(+) condition revealed no significant difference when compared to A1405 Fe(+) (data not shown). These findings are consistent with our proteomic analysis, supporting the conclusion that both iron depletion and MpkA deletion synergistically enhance siderophore biosynthesis in *A. nidulans*.

**Figure 6.**
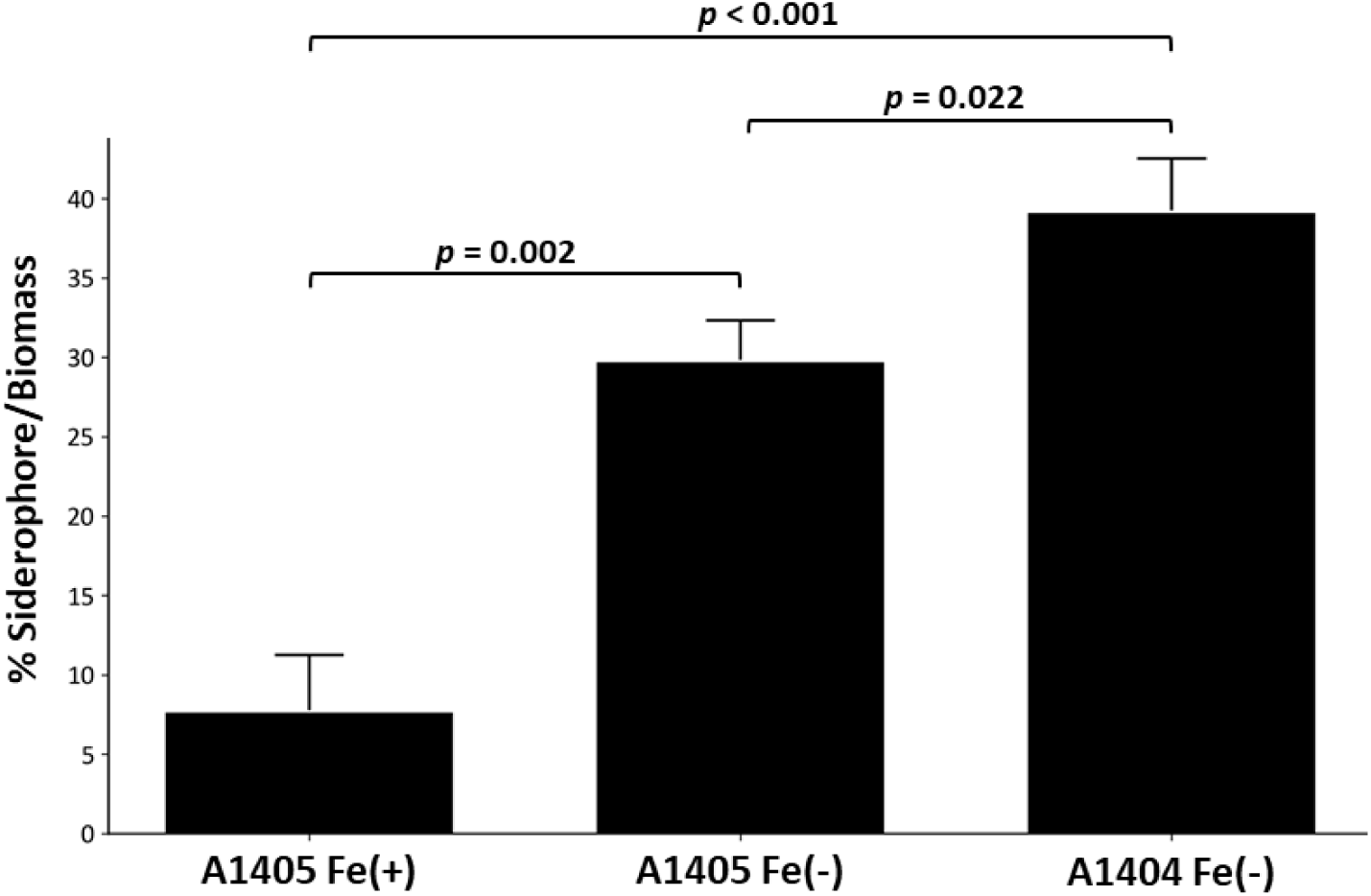
Siderophore production normalized to biomass, under iron limitation and MpkA deficiency, after 20 hours of growth in minimal medium. Iron-depleted conditions were established by culturing in minimal medium without iron supplementation. Siderophore levels were quantified using the CAS assay and expressed as a percentage. Biomass was measured as dry cell weight (mg/g).

## Conclusion

While fungal virulence is closely linked to its ability to compete with the host for iron acquisition, the MAP kinase MpkA may influence this process by modulating the siderophore system. Consequently, comprehensive analysis of proteome expression and phosphorylation dynamics is essential for understanding fungal adaptation. This study demonstrates the application of DIA-PASEF for high-throughput proteome and phospho-proteome profiling in *A. nidulans*. Our workflow enabled deep proteome coverage using short chromatographic gradients and revealed that the *mpkA* deletion, particularly under iron-limiting conditions, leads to extensive remodeling of the integrated proteomic landscape, encompassing both protein expression and phosphorylation networks.

Gene Ontology (GO) terms related to ATP and ADP metabolism were uniquely enriched under iron-deficient conditions, indicating increased energy cycling activity in the *mpkA*-deleted strain. This alteration in the ATP and ADP pool may be associated with elevated PPP flux ^80, 81^. In addition, GO terms related to secondary metabolic processes indicated potential upregulation of the tricarboxylic acid (TCA) cycle in the Δ*mpkA* strain under iron-limiting conditions, which could supply key precursors for siderophore biosynthesis and contributing to the observed elevation in siderophore production. Additionally, GO terms related to transport, secretion, and reproduction were specifically enriched among differentially phosphorylated proteins (DPPs) under iron-deficient conditions in the *mpkA* mutant, suggesting potential effects on host infection mechanisms. Together, these findings provide new insights into fungal signaling and adaptive responses to environmental and genetic perturbations.

## ASSOCIATED CONTENT

### Supporting Information

**Supplementary Table S1.** Assignment of pooled eluates.

**Supplementary Table S2.** List of proteins identified in A1405 cultured in YGV medium using DDA-PASEF and DIA-PASEF approaches.

**Supplementary Table S3.** List of proteins and phospho-sites identified in A1405 and A1404 cultured in minimal media with and without iron using DIA-PASEF approaches.

**Supplementary Table S4.** Gene Ontology biological process terms enriched among DEPs and DPPs in the A1404 vs A1405 comparison under iron-replete and iron-depleted conditions.

**Supplementary Table S5.** Gene Ontology biological process terms enriched among DEPs and DPPs in the A1404 vs A1405 comparison were particularly prominent under iron-depleted conditions.

## AUTHOR INFORMATION

### Authors

JungHun Lee - Department of Chemical, Biochemical, and Environmental Engineering, University of Maryland Baltimore County, Baltimore, MD 21250, United States

Olivia K. West - Department of Chemical, Biochemical, and Environmental Engineering, University of Maryland Baltimore County, Baltimore, MD 21250, United Statesd

Walker D. Huso - Department of Chemical, Biochemical, and Environmental Engineering, University of Maryland Baltimore County, Baltimore, MD 21250, United States

Alexander G. Doan - Department of Chemical, Biochemical, and Environmental Engineering, University of Maryland Baltimore County, Baltimore, MD 21250, United States

Kelsey J. Grey - Department of Chemical, Biochemical, and Environmental Engineering, University of Maryland Baltimore County, Baltimore, MD 21250, United States

Harley Edwards - Department of Chemical, Biochemical, and Environmental Engineering, University of Maryland Baltimore County, Baltimore, MD 21250, United States

Mark R. Marten - Department of Chemical, Biochemical, and Environmental Engineering, University of Maryland Baltimore County, Baltimore, MD 21250, United States

Jasmine T. Tran - Department of Chemical and Biomolecular Engineering, Johns Hopkins University, Baltimore, MD 21218, United States

Dylan R. Carmen - Department of Chemical and Biomolecular Engineering, Johns Hopkins University, Baltimore, MD 21218, United States

Michael J. Betenbaugh - Department of Chemical and Biomolecular Engineering, Johns Hopkins University, Baltimore, MD 21218, United States

Ranjan Srivastava - Department of Chemical and Biomolecular Engineering, University of Connecticut, Storrs, CT 06269, United States

Steven D. Harris - Iowa State University, Department of Plant Pathology, Entomology and Microbiology, Ames, IA 50011, United States

### Author Contributions

JH.L. designed the experiments. JH.L. and O.K.W. did strain culture. JH.L. performed protein and siderophore sample preparation and analysis. M.R.M. supervised the project. and provided funding. JH.L. wrote the original draft. JH.L., O.K.W., W.D.H., A.G.D., K.J.G., H.E., J.T.T., D.R.C., M.J.B., R.S., S.D.H., and M.R.M. assisted in developing concepts, understanding results and revising the manuscript.

### Notes

The authors declare no competing financial interest.

## ACKNOWLEDGMENT

Peptide samples were analyzed using a timsTOF Pro2 mass spectrometer at the Molecular Characterization and Analysis Complex (MCAC), University of Maryland, Baltimore County (UMBC).

## ABBREVIATIONS

QTOF: quadrupole time-of-flight;
timsTOF: trapped ion mobility time of flight;
PASEF: parallel Accumulation–Serial Fragmentation;
DDA: data-dependent acquisition
DIA: data-independent acquisition;
DIA-PASEF: data-independent acquisition–parallel accumulation serial fragmentation;
DDA-PASEF: data-dependent acquisition–parallel accumulation serial fragmentation;
CWI: cell wall integrity;
MpkA: mitogen-activated protein kinase A;
DEPs: differentially expressed proteins;
DPPs: differentially phosphorylated proteins;
GO: gene ontology

